# Cellular polarity asymmetrically functionalizes pathogen recognition receptor-mediated intrinsic immune response in human intestinal epithelium cells

**DOI:** 10.1101/450668

**Authors:** Megan L. Stanifer, Stephanie Muenchau, Markus Mukenhirn, Kalliopi Pervolaraki, Takashi Kanaya, Dorothee Albrecht, Charlotte Odendall, Thomas Hielscher, Volker Haucke, Jonathan Kagan, Sina Bartfeld, Hiroshi Ohno, Steeve Boulant

## Abstract

Intestinal epithelial cells (IECs) act as a physical barrier separating the commensal-containing intestinal tract from the sterile interior. These cells have found a complex balance allowing them to be prepared for pathogen attacks while still tolerating the presence of bacteria/viral stimuli present in the lumen of the gut. Using primary human IECs, we probed the mechanisms, which allow for such a tolerance. We discovered that viral infection emanating from the basolateral side of IECs elicited a stronger intrinsic immune response as compared to lumenal apical infection. We determined that this asymmetric immune response was driven by the clathrin-sorting adapter AP-1B which mediates the polarized sorting of Toll-like receptor 3 (TLR3) toward the basolateral side of IECs. Mice and human IECs lacking AP-1B showed an exacerbated immune response following apical stimulation. Together these results suggest a model where the cellular polarity program plays an integral role in the ability of IECs to partially tolerate apical commensals while remaining fully responsive against invasive basolateral pathogens.

## Introduction

Intestinal epithelial cells (IECs) lining the gastrointestinal tract constitute the primary barrier separating us from the outside environment. These cells protect the human body from enteric bacterial and viral pathogens (Peterson and Artis, 2014). They sense and combat these pathogens using Pathogen Recognition Receptors (PRRs) (e.g. the Toll-like receptors (TLRs)) to trigger an intrinsic innate immune response (Pott and Hornef, 2012, Fukata and Arditi, 2013, Arpaia and Barton, 2011, Barton and Medzhitov, 2003). Importantly, IECs are in constant contact with the ever-present microbiota and therefore have developed mechanisms to tolerate the presence of the commensals while maintaining responsiveness against pathogen (Fukata and Arditi, 2013). The intrinsic innate immune response is regulated not only by complex signal transduction pathways downstream of PRRs but also by compartmentalization of these receptors (Odendall and Kagan, 2017, Chow et al., 2015). This compartmentalization is described to be an important mechanism by which cells avoid self-recognition (Yu and Gao, 2015, Kagan and Barton, 2014). In IECs, this compartmentalization of innate immune functions is believed to be even more critical to maintain gut homeostasis (Yu and Gao, 2015). IECs are polarized, they display a unique apical membrane facing the lumen of the gut and a basolateral membrane facing the lamina propria (Yu and Gao, 2015, Rodriguez-Boulan and Macara, 2014). Enteric pathogens and commensals are normally located in the lumen of the gut thereby generally challenging hIECs through their apical membrane. However, when the barrier function is lost, in the presence of invasive pathogens, or during spreading of infection, pathogens and commensals can gain access the basolateral side of hIECs. The concept that PRRs can induce a different signaling depending on whether the pathogen challenges emanate from the apical or basolateral side of hIECs have been pioneered by studying TLR9 and TLR5 (Gewirtz et al., 2001, Lee et al., 2006). The mechanisms leading to this polarized immune response have been described to be mediated by a different signal transduction pathway between apical or basolateral stimulation for TLR9 (Lee et al., 2006) and due to the asymmetric distribution of the PRRs in hIECs for TLR5 (Gewirtz et al., 2001). The precise intracellular localization of TLRs and, most importantly, which ones are actually expressed in IECs has been a matter of debate (Price et al., 2018). Additionally, how viruses are detected and combatted in human IECs has been largely understudied. It remains unknown whether virus specific PRRs (i.e. TLR3) can be asymmetrically distributed thereby providing IECs the capacity to distinguish between viral infection emanating from the apical or their basolateral side and as a consequence mount a different immune response.

In our study, we determined that IECs respond to viral infection in a side specific manner. We could directly correlate this polarized immune response to the asymmetric distribution of TLR3 towards the basolateral side of IECs. We identify the clathrin sorting adaptor protein AP-1 as the key machinery to sort TLR3 to the basolateral side of IECs. Our results suggest a mechanism used by IECs to generate a moderate immune response against stimuli emanating from the physiological lumenal side while remaining fully responsive against invasive pathogens or in conditions of barrier integrity loss where stimuli can access the normally sterile basolateral side.

## Results

To address the extent by which TLRs act in a polarized manner in intestinal epithelial cells, the model human IEC line T84 cells were polarized on transwells (Madara et al., 1987). Cellular polarization was controlled through immunostaining of the tight junction belt (data not shown and (Stanifer et al., 2016)). The integrity of the epithelial barrier, a pre-requisite for side specific stimulation, was validated by monitoring the Trans-Epithelium Electrical Resistance (TEER) (Sup. Figure 1A) and by controlling the capacity of the hIEC monolayer to block dextran diffusion in a transwell diffusion assay (Sup. Figure 1B). These polarized hIECs were used to evaluate the relative expression of each TLR by q-RT-PCR. Results showed that TLR2, 3, 4 and 5 were the most expressed in hIECs (Figure 1A). Following side-specific stimulations by TLR agonists (Figure 1B), the specific response of hIECs was evaluated by monitoring the transcription upregulation of both IL-6 and type III interferon (IFNλ2/3). Stimulation with TLR7, 8 and 9 agonists did not induce an immune response in hIECs which was consistent with their low expression levels (Figure 1A-D). However, these agonists were fully potent in inducing IL-6 production in pBMCs (Sup. Figure 2A). Importantly, while treatments of the cells with TLR2, 3, 4 and 5 agonists resulted in the production of IL-6 (and IL-8, data not shown) and IFNλ, basolateral stimulation of the cells by TLR3 agonist induced a significantly higher production of IFNλ compared to apical stimulation (Figure 1C-D). This stronger induction of interferon upon basolateral stimulation with the TLR3 agonist was true at any time post-treatment (Figure 1E). This TLR3-specific polarized immune response was not the result of a more efficient update of the agonist from the basolateral side of the cells as quantification revealed that the kinetics of fluorescently labeled TLR3 agonist uptake was identical between the apical and basolateral side of hIECs (Sup. Figure 2B). Together, these results suggest that hIECs can distinguish between apical and basolateral stimulation of TLR3 and adapt their response in a side specific manner.

**Figure 1.**
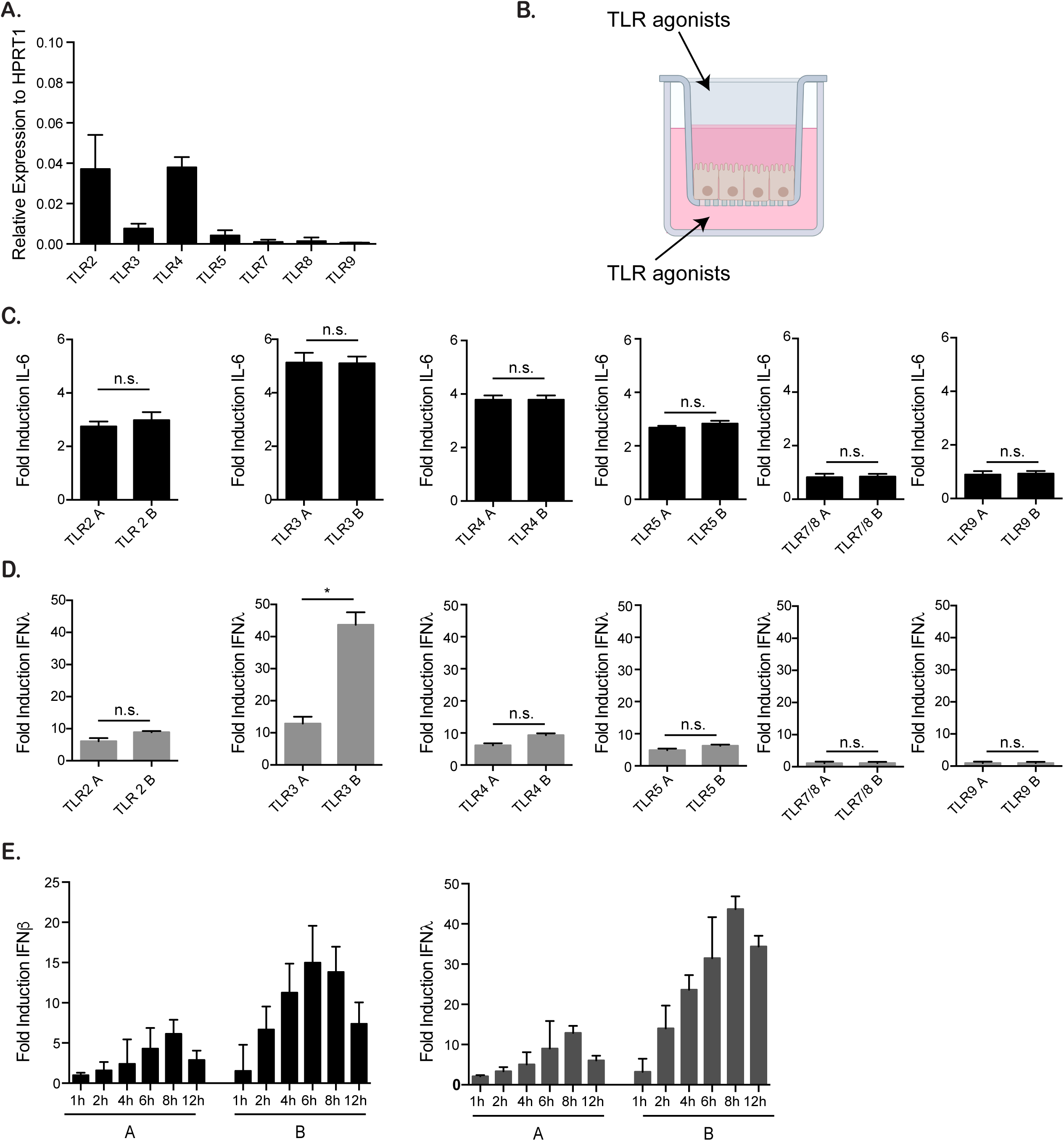
T84 cells respond to TLR3 agonist in a side-specific manner. (A). The relative expression of TLR2-9 was evaluated in T84 cells by q-RT-PCR. Results are expressed as a relative expression to the house-keeping gene HPRT1. (B) Diagram depicting the side specific stimulation (apical vs. basolateral) of polarized T84 cells grown on transwells by TLR agonists. (C) T84 cells were seeded on transwells and after establishment of barrier function, cells were treated with the following TLR agonists: TLR1/2 (PAM3CSK4, 1 μg/mL); TLR3 (1:1 HMW and LMW poly I:C, 10 μg/mL); TLR4 (LPS-EK, 0.01 μg/mL); TLR5 (Flagellin-ST, 1 μg/mL); TLR7/8 (R848, 1 μg/mL), TLR9 (ODN 2395 5 μM). 6 hours post-treatment, RNA was harvested and evaluated for the up-regulation of IL-6 by q-RT-PCR. Results are normalized to mock-treated samples. (D) Same as C except type III IFN (IFNλ2/3) was measured. (E) T84 cells were seeded on transwells and treated with 10 μg/mL of poly I:C. Cells were harvested at the indicated time points and up-regulation of both type I (IFNβ) and type III (IFNλ2/3) IFNs were analyzed by q-RT-PCR. Experiments were performed in triplicate, error bars indicate the standard deviation. ns=not significant, *<P.05 (unpaired t-test)

### Basolateral viral infection of hIECs induces a stronger immune response compared to apical infection

To control that the polarized TLR3 immune response was not agonist specific we infected hIECs with mammalian reovirus which is actively sensed by TLR3 during infection (Stanifer et al., 2016). Polarized hIECs grown on transwells were infected in a polarized manner (Figure 2A) and virus replication and cellular immune response was followed overtime as function of infection side (apical vs. basolateral). Results showed that the onset of viral replication was identical between apical and basolateral infection. No significant differences between apical and basolateral infection could be detected in the efficacy of virus uptake (Sup. Figure 3A), in the number of cells infected during the primary round of infection (Figure 2B-C), and in viral replication as measured by viral genome load and production of the viral non-structural protein μNS overtime (Sup. Figure 3B-C). However, at late time points and during multiple rounds of infection, apical infection resulted in a higher level of replication and production of progeny viruses (Figure 2D-E and Sup. Figure 3C-E). Interestingly, the lower production of *de novo* viruses following basolateral infection correlated with a stronger intrinsic innate immune response at both the RNA and protein level (Figure 2F-H). A similar stronger immune response following basolateral infection was also observed when using a different hIEC cell line (SKCO15) (Le Bivic et al., 1989) (Sup. Figure 4). Additionally, as observed using the TLR3 agonist poly I:C (Figure 1E), basolateral viral infection of hIECs resulted in a higher production of interferon at all times post-infection at both the RNA and protein levels (Sup. Figure 5).

**Figure 2.**
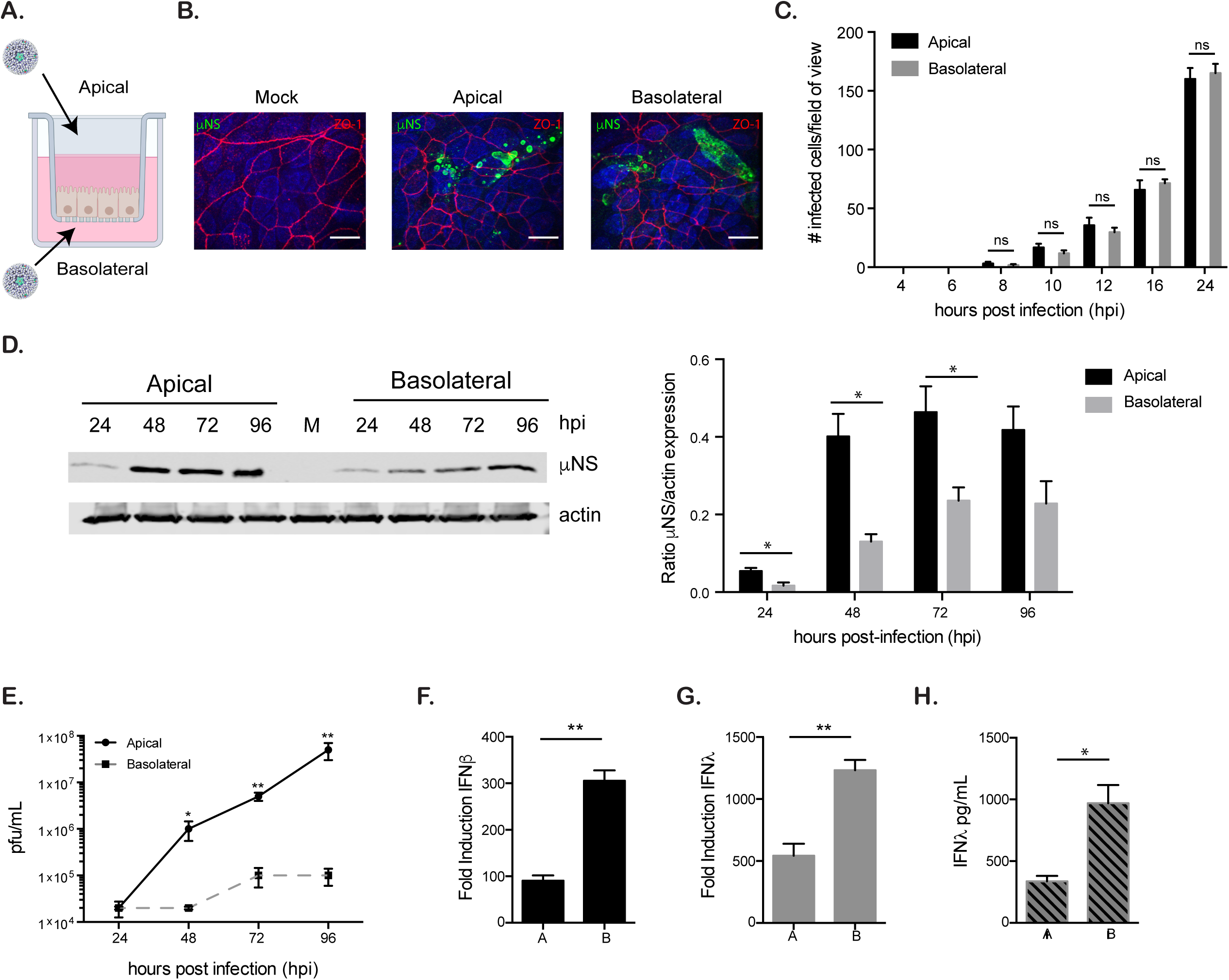
Intrinsic immune response of T84 cells to viral infection is side specific. T84 cells were seeded onto transwell inserts and allowed to establish barrier function. (A) Schematic depicting infection of T84 cells in a side specific manner. (B) Polarized T84 cells were infected apically or basolaterally with mammalian reovirus (MRV). 16 hours post-infection (hpi) cells were fixed and indirect immunofluorescence was performed to control the integrity of the tight junction belt (tight junction protein ZO-1, red) and to evaluate the viral non-structural protein μNS (green). Nuclei were visualized with DAPI. (C) Polarized T84 cells were infected apically or basolaterally with MRV. At indicated time points, cells were fixed and stained for the viral non-structural protein μNS. The number of infected cells was counted in ten independent fields of view for each sample. (D) Polarized T84 cells were infected apically or basolaterally with MRV. At indicated time points cells were harvested and Western blot was used to detect the amount of the non-structural protein μNS in each sample (left panel). The relative amount of μNS normalized to actin was quantified for each time point post-infection (right panel). (E) Same as D except cells were lysed and production of progeny viruses was evaluated using a plaque assay. (F-G) Polarized T84 cells were infected apically or basolaterally with MRV. 16 hpi, cells were harvested and the production of interferons was analyzed by q-RT-PCR. (F) Type I (IFNβ). (G) Type III (IFNλ2/3). (H) Same as G except that production of type III (IFNλ2/3) IFN was analyzed by ELISA. Experiments were performed in triplicate, representative images for Western blot and immunofluorescence are shown, error bars indicate the standard deviation. ns=not significant, *<P.05, **P < 0.01 (unpaired t-test)

This higher production of interferon following basolateral stimulation and infection could be due to an increased RNA stability or sustained transcription. To address this possibility, cells were treated with actinomycin to inhibit new RNA synthesis (Sup. Figure 6A). The half-life of both apical and basolateral IFNλ2/3 transcripts were found to be similar, however basolateral infection lead to a slightly lower half-life of IFNβ1 transcripts compared to apical infection (Sup. Figure 6B-C). These finding argue that the stronger immune response generated upon basolateral challenges is not the result of differences in the stability of the IFN transcripts. To gain a global understanding of the extent by which apical and basolateral infection/stimulation leads to a different immune response hIECs were infected in side specific manner and their transcriptional profiles was analyzed by microarray. Results show that infection from both sides leads to an increase in immune response related genes however, basolateral infection resulted in a greater number of upregulated genes as well as a higher induction of the genes also upregulated upon apical infection (Sup. Figure 7). In addition, KEGG analysis shows that the upregulated genes are predominantly associated with immune response pathways and interferon-mediated responses. (Sup. Figure 7C).

All together these data show that hIECs can react in a side specific manner to TLR3 agonist and viral infection. Basolateral infection/stimulation induces a stronger interferon-mediated immune response compared to the same stimuli coming from their apical side.

### Primary intestinal cells respond in a side specific manner to TLR3 stimuli

To validate our findings in human primary cells we exploited human mini-gut organoids. To perform a side specific stimulation/infection, organoids were grown and differentiated on transwells (Figure 3A). Cell differentiation and the integrity of the barrier function was controlled by immunostaining and TEER measurements (Figure 3B and data not shown). Primary hIECs were infected from either their apical or basolateral side. Although infectivity was not side dependent, basolateral infection lead to a stronger production of interferon (Figure 3C-E). Similar findings were observed when performing side specific infection of 3D organoids. Using microinjection, MRV was introduced within the lumen of the organoids to allow for apical infection or juxtaposed at the organoid periphery to promote basolateral infection (Sup Figure 8A). Following validation that our apical and basolateral microinjection approaches led to equivalent infection of the human mini-gut organoids (Sup Figure 8B-C), we confirmed that basolateral infection lead to a stronger production of interferon compared to apical infection (Sup Figure 8D). To investigate whether this polarized immune response also applies to other TLRs, we performed a side specific stimulation with TLR agonist of primary human organoids on transwells. As observed in our T84 cell line, primary hIECs mostly express TLR2, 3, 4 and 5 (Sup. Figure 8E) which was consistent with previous work using murine IECs (Price et al., 2018). Importantly, although these cells responded to the respective agonists (Figure 3F-G), they only show a polarized interferon induction upon TLR3 stimulation (Figure 3G). Altogether, our results demonstrate that hIECs respond in an asymmetric manner to TLR3 stimulation.

**Figure 3.**
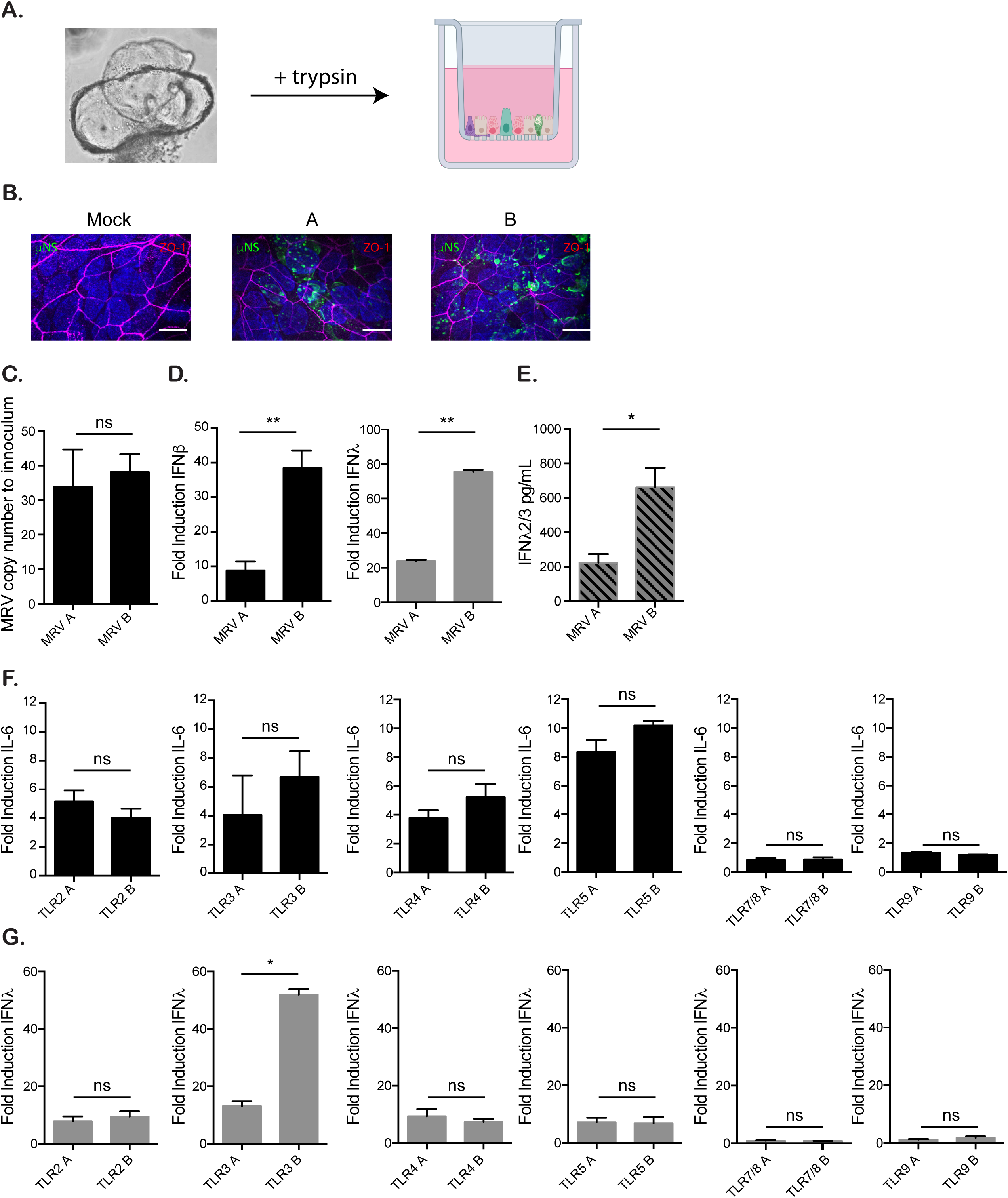
Human organoids respond in a polarized manner to TLR3 agonist. (A) Human organoids were seeded onto transwells and allowed to polarize and differentiate for five days (see method for details). Polarized monolayers were infected with MRV apically or basolaterally and assessed 16 hpi for (B) viral infection by indirect immunofluorescence (green=μNS, red=ZO-1, blue=DAPI) and (C) for the amount of viral genome present. (D) The production of both type I (IFNβ) and type III (IFNλ2/3) interferons was evaluated by q-RT-PCR. (E) Same as D except type III IFN secretion was quantified by ELISA. (F) Colon organoids were seeded on transwells and treated with the indicated TLR agonist. TLR1/2 (PAM3CSK4, 1 μg/mL); TLR3 (1:1 HMW and LMW poly I:C, 10 μg/mL); TLR4 (LPS-EK, 0.01 μg/mL); TLR5 (Flagellin-ST, 1 μg/mL); TLR7/8 (R848, 1 μg/mL), TLR9 (ODN 2395 5 μM). 6 hours post-treatment RNA was harvested and evaluated for the up-regulation of IL-6 by q-RT-PCR. Results are normalized to mock-treated samples. (G) Same as F, except type III IFN (IFNλ2/3) was evaluated. All experiments were performed in triplicate. Representative immunofluorescence images are shown. Error bars indicate the standard deviation. Scale bar in (B) represents 10 μm. ns=not significant, *<P.05, **P < 0.01 (unpaired t-test).

### The clathrin sorting adapter AP-1B mediates the basolateral localization of TLR3 in hIECs

To address whether the polarized response to infection/stimulation was due to a polarized distribution of the pathogen recognition receptor, we performed immunostaining of TLR3 in hIECs. Human organoids were seeded as a 2D monolayer as described above (Figure 3A), and results show that TLR3 was restricted to the basolateral membrane of hIECs (Figure 4A). Similar results were obtained with both T84 cells grown as spheroids and primary hIECs grown as 3D organoids (Figure 4B-C). Quantification revealed that TLR3 was exclusively localized to the basolateral membrane in human organoids and was mostly localized to the basolateral membrane in T84 cells (Figure 4D).

**Figure 4.**
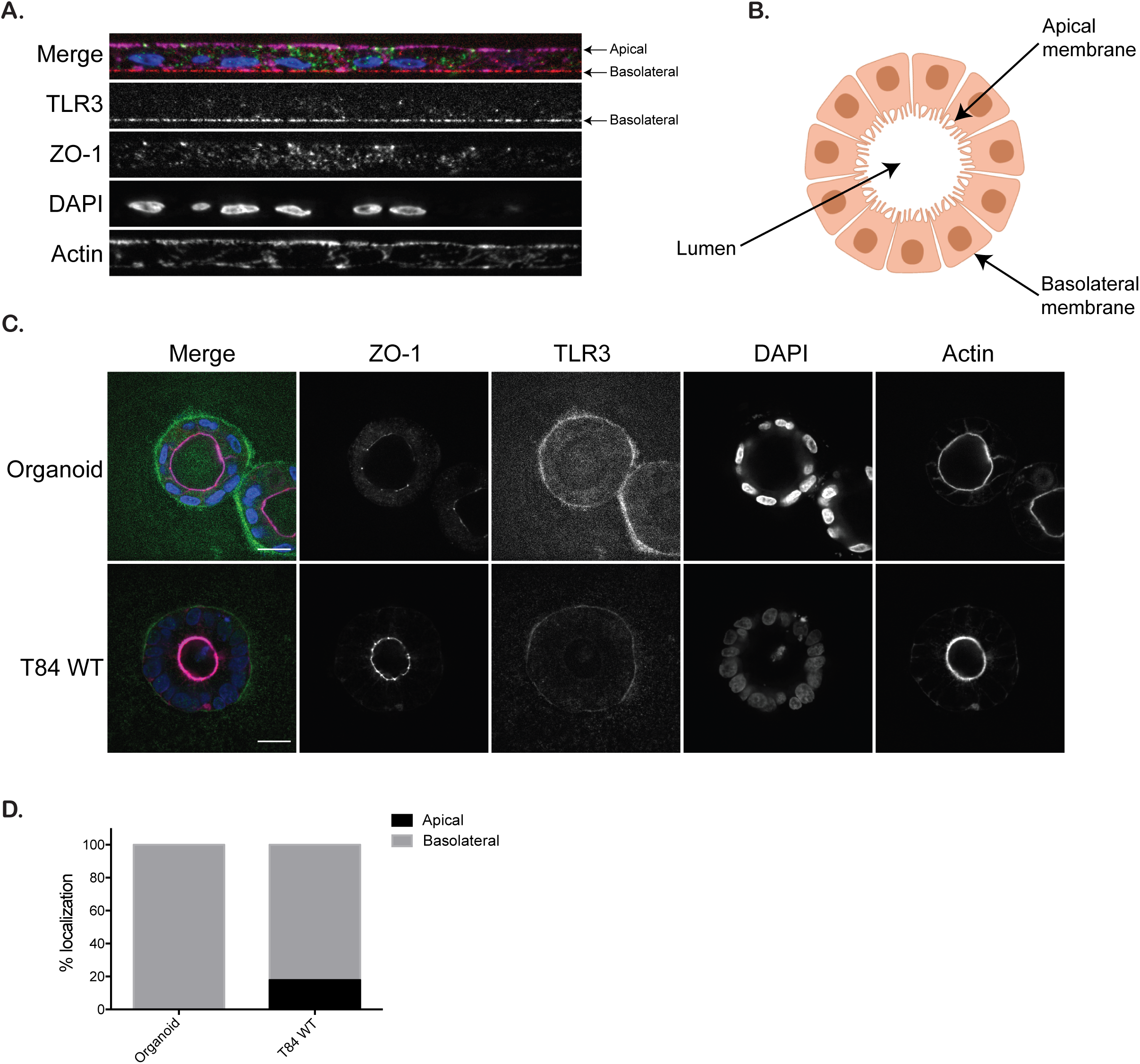
TLR3 is localized to the basolateral side of intestinal epithelial cells. (A). Human colon organoids were seeded flat on a Labtek chamber pre-coated with Matrigel. Following fixation with PFA, cells were immunostained against TLR3 (red), ZO-1 (green), actin was labeled with phalloidin 647 (magenta) and the nuclei were stained with DAPI (blue). (B) Schematic depicting a T84-derived spheroid or a small organoid, with the apical membrane of hIECs facing the lumen of the structure and their basolateral membrane facing outward. (C) Human colon organoids and T84-derived spheroids were immunostained against TLR3 (green), ZO-1 (red), actin was labeled with phalloidin 647 (magenta) and the nuclei were stained with DAPI (blue). (D) The percentage of spheroids displaying TLR3 at the apical or basolateral membrane of hIECs was quantified (For organoids n=51, for T84 cells n=84). Experiments were performed in triplicate, representative images are shown. Scale bar represents 20 μm.

The clathrin sorting adapter AP-1 is well known for sorting proteins to the basolateral membrane of polarized cells ((Folsch, 2015). AP-1 comes in two forms; the ubiquitously expressed AP-1A and the epithelial specific AP-1B. To investigate if AP-1B was responsible for both the polarized distribution of TLR3 and the polarized immune response, we used shRNA to knock-down AP-1B in our T84 cell line model (Sup. Figure 9A). Immunostaining of TLR3 revealed that upon AP-1B down regulation, TLR3 was re-localized from the basolateral membrane to the apical side of hIECs (Figure 5A-C). Quantification revealed that most of the AP-1B knock-down spheroids display TLR3 at their apical side (Figure 5D). After controlling the barrier function and integrity of AP-1B knock-down hIECs (Sup. Figure 9B-D), WT and AP-1B knock-down cells were infected in a side specific manner with reovirus. As observed before, basolateral infection of WT hIECs resulted in a stronger production of interferon compared to apical infection. Interestingly, knock-down of AP-1B resulted in an inversion of this polarized response leading to a stronger production of interferon upon apical challenges both at the transcript (Figure 6A) and protein level (Sup. Figure 9E). This stronger interferon production upon apical infection of AP-1B knock-down cells fully correlates with the observation that TLR3 is re-localized to the apical side of hIECs (Figure 5A-D). To control that AP-1B knock-down did not impact global TLR mediated signaling, WT and AP-1B knock-down cells were treated with TLR agonists in a side specific manner. No significant differences were observed for TLR2, 4, 5, 7/8-, and 9-mediated signaling upon AP-1B knock-down (Sup. Figure 10). Importantly, similar to viral infection, an inversion of polarized immune response in AP-1B knock-down cells was observed leading to a higher induction of interferon upon apical treatment of the cells with the TLR3 agonist (Figure 6B and Sup. Figure 10B). To validate that the observed increased immune response in AP-1B knock-down cells upon apical challenges was directly mediated by TLR3, we used a pharmacological inhibitor approach. As TLR3 requires TRIF as adapter protein to mediate downstream signaling (Yamamoto et al., 2003), we used a TRIF specific inhibitor. Results confirm that the observed immune response generated by hIECs upon viral infection is largely due to TLR3 (Sup. Figure 11A). Importantly, inhibition of TLR3-mediated signaling in AP-1B knock-down cells resulted in a strong reduction of immune response upon apical infection (Sup. Figure 11B). These results strongly suggest that the observed inversion of polarized immune response in AP-1B knock-down is directly associated with the re-localization of TLR3 to the apical side of hIECs. All together these data support a model where the clathrin sorting adapter AP-1 mediates the polarized sorting of TLR3 in hEICs rendering them more responsive to pathogen challenges emanating from their basolateral side.

**Figure 5.**
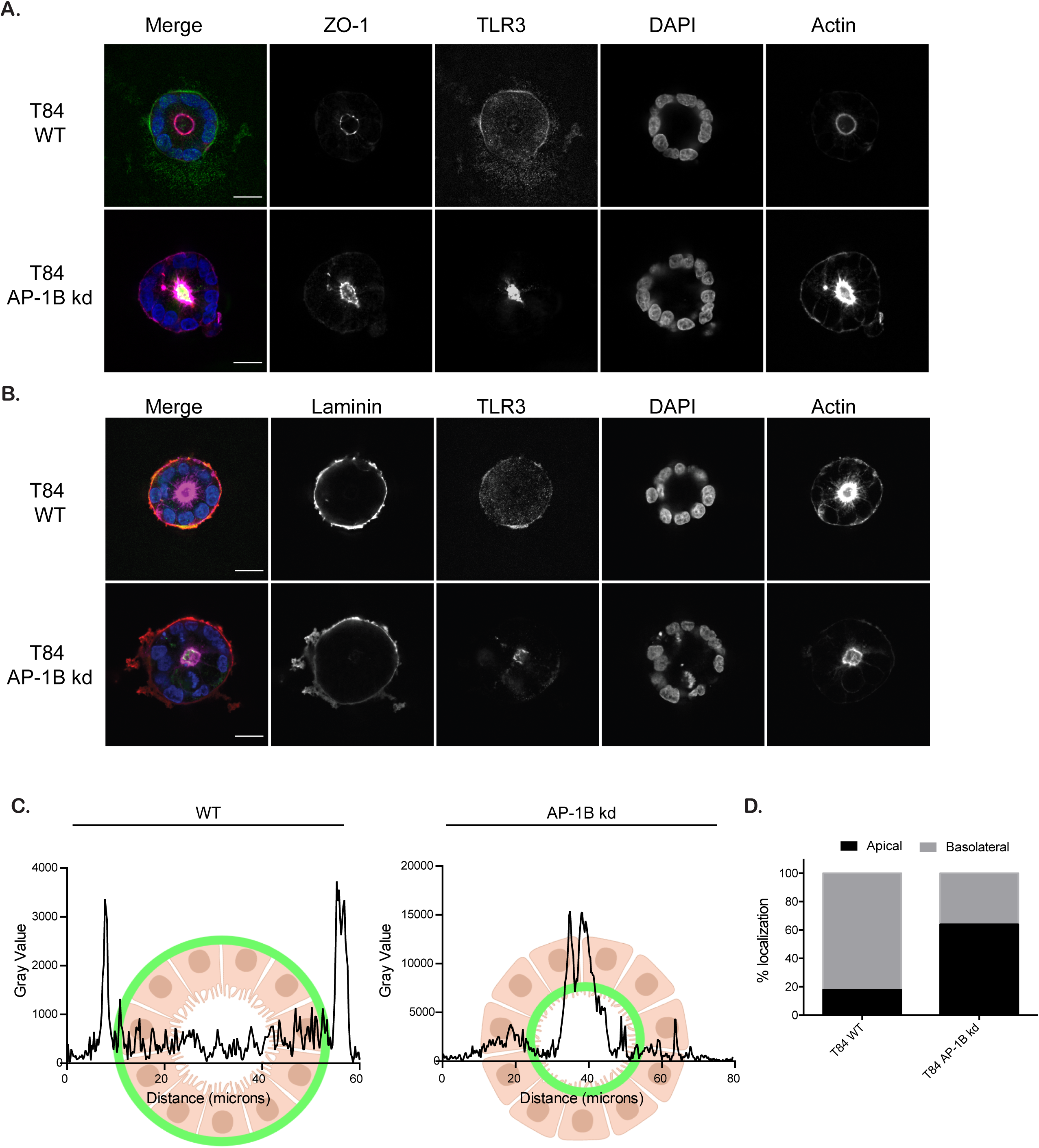
The clathrin sorting adaptor AP-1B is responsible for the basolateral localization of TLR3 in hIECs. (A) WT and AP-1B knock-down T84 spheroids were immunostained against TLR3 (green), ZO-1 (red), actin was labeled with phalloidin 647 (magenta) and the nuclei were stained with DAPI (blue). (B) Same as A except cells were immunostained for Laminin (red) as a basolateral marker. (C) Quantification of the localization of TLR3 in WT and AP-1B knock-down T84 spheroids. The relative fluorescence intensity of TLR3 is depicted along a transversal line going in the middle of the spheroids. (D) The percentage of spheroids displaying TLR3 at the apical or basolateral membrane of hIECs was quantified. (for WT spheroids n=84, for AP-1B knock-down T84 spheroids n=111). Experiments were performed in triplicate, representative images are shown. Scale bar represents 20 μm.

**Figure 6.**
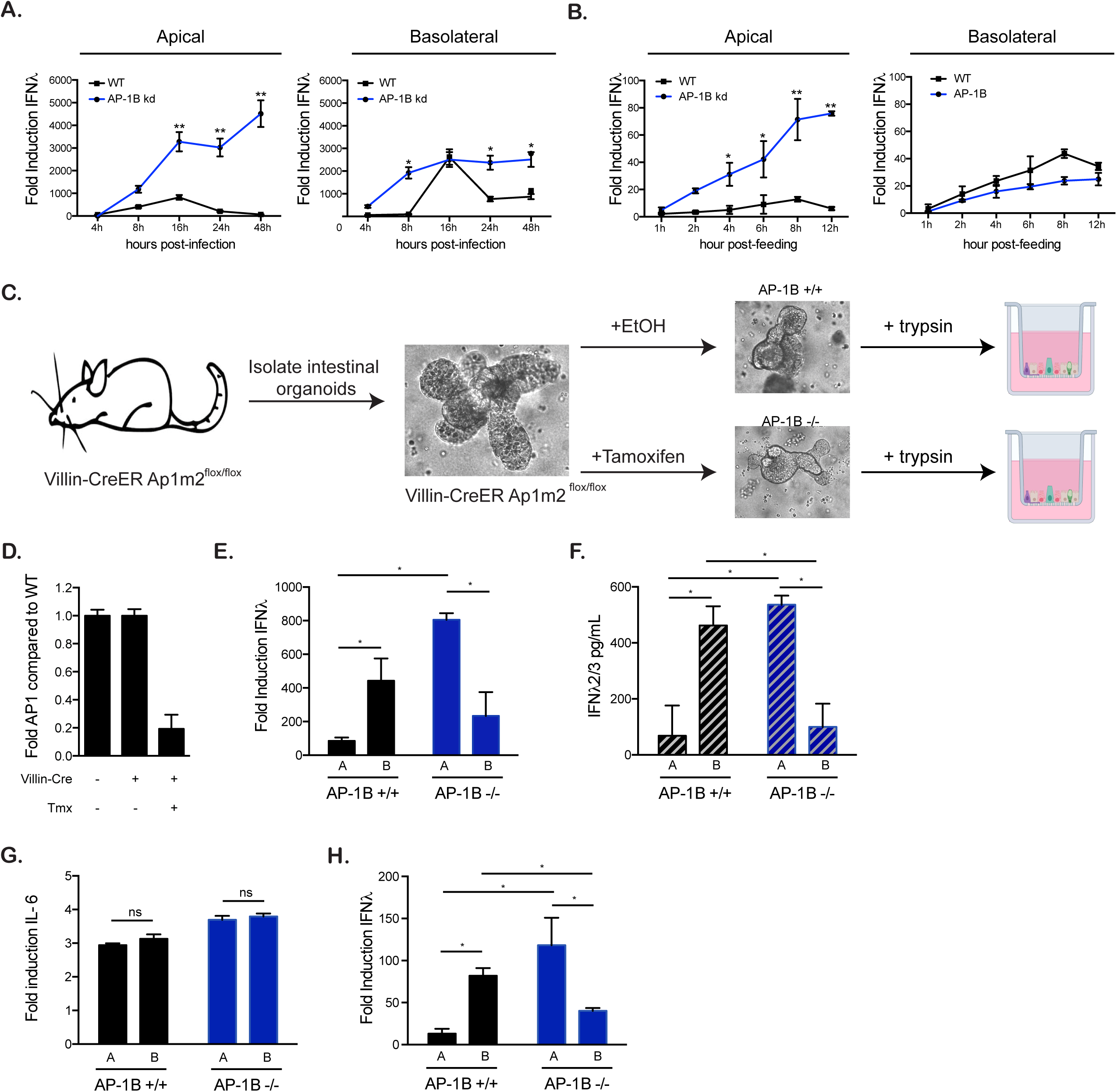
The clathrin sorting adaptor AP-1B is responsible for the TLR3-mediated polarized immune response. (A) WT and AP-1B knock-down T84 cells were polarized on transwell inserts. Cells were infected with MRV apically or basolaterally. At indicated time points, the production of type III (IFNλ2/3) interferon was evaluated by q-RT-PCR. (B) Same as A except polarized T84 cells were stimulated with the TLR3 agonist poly I:C (10 μg/mL). (C) Schematic for the generation of the inducible murine AP-1B knock-out organoids from the Villin-CreER Ap1m2 ^flox/flox^ mice. (D). AP-1B knock-out was confirmed by q-RT-PCR for exon 4-5 of the Ap1m2 locus. (E) Non-tamoxifen treated Villin-CreER Ap1m2 ^flox/flox^ mice (AP-1B+/+) and tamoxifen treated Villin-CreER Ap1m2 ^flox/flox^ mice (AP-1B -/-) derived organoids were seeded onto transwells and infected with MRV in a polarized manner. 48 hpi, cells were harvested and analyzed for the induction of IFNλ2/3 by q-RT-PCR. (F) Same as E, except IFNλ2/3 was analyzed by ELISA. (G) Non-tamoxifen treated Villin-CreER Ap1m2 ^flox/flox^ mice (AP-1B+/+) and tamoxifen treated Villin-CreER Ap1m2 ^flox/flox^ mice (AP-1B -/-) derived organoids were seeded onto transwells and stimulated with the TLR3 agonist poly I:C (10 μg/mL). 8 hours post-stimulation, cells were harvested and analyzed for the induction of IL-6 by q-RT-PCR (H) Same as G, except IFNλ2/3 was analyzed. All experiments were performed in triplicate. Error bars indicate the standard deviation. ns=not significant, *<P.05, **P < 0.01 (unpaired t-test).

### IECs derived from AP-1B knock-out mice show a reversion of their polarized immune response

To confirm that AP-1B was contributing to the observed polarized immune response we evaluated mice, which express an inducible intestinal specific knock-out of AP-1B (villin-CreER Ap1m2^flox/flox^, (Takahashi et al., 2011)) (Figure 6C). Intestinal organoids were generated from these mice and exploited for apical and basolateral infection as side specific infection is not possible *in vivo*. While apical infection can be mimicked through gavaging of virus, basolateral infection cannot be controlled as intraperitoneal infection will also deliver virus to the surrounding immune cells and stroma cells thereby resulting in an immune response generated not only by IECs but also by other cell types. Organoids generated from villin-CreER Ap1m2^flox/flox^ were treated with or without tamoxifen and the loss of AP-1B was controlled by qPCR (Figure 6C-D) (see method). AP-1B^+/+^ and AP-1B^-/-^ organoids were seeded onto transwells and infected apically or basolaterally with reovirus. The innate immune induction was controlled by q-RT-PCR and confirmed that similar to our T84 cell model and to our human organoids, murine organoids also show a polarized immune response where a basolateral infection leads to a stronger production of IFN compared to an apical one at both the RNA and protein level (Figure 6E-F). Importantly, in AP-1B knock-out murine organoids, an inversion of polarized immune response was observed where apical infection lead to a higher production of interferon compared to basolateral infection (Figure 6E-F). Similar findings were observed when using a TLR3 agonist to induce an immune response in murine organoids further confirming the role of AP-1B in TLR3-mediated polarized interferon induction (Figure 6G-H).

## Discussion

In this study we unraveled that the extent of the immune response which is generated downstream TLR3 depends on the side (apical vs. basolateral) of infection/stimulation. Precisely, we show that basolateral infection/stimulation leads to a higher induction and secretion of IFNλ2/3 compared to an apical infection. We show that the clathrin adaptor AP-1B sorting machinery allows for the asymmetric distribution of TLR3, allowing IECs to be more responsive from their basolateral side. Interference with AP-1B renders IECs more sensitive to apical stimuli. We propose that this polarized immune response represents a unique immune-strategy to create a homeostatic interface between the complex microbial environment of the gut lumen and the intestinal epithelial barrier. It allows IECs to restrain their immune response against the naturally present apical stimuli while remaining fully responsive to infection/stimulation emanating from the physiologically sterile basolateral side, which will only be accessible during loss of barrier function or during infection by invasive pathogens.

Only a few studies have tried to correlate the specific localization of TLRs at the apical and basolateral side of IECs with their capacity to respond to PAMPs. TLR5 has been shown to be located on the basolateral membrane, which was suggested to be critical to limit overstimulation of IECs by the commensal bacterial components flagellin (Gewirtz et al., 2001). TLR2 and 4 have been described to be redistributed near the basolateral side of IECs upon activation (Cario et al., 2002, Hiemstra et al., 2015). It was suggested that this trafficking is a mechanism used by IECs to transmit information from the lumen into underlying the immune cells (Hiemstra et al., 2015). TLR9 was shown to be located in both the apical and basolateral membranes but has been described to signal in a polarized manner. Basolateral stimulation of TLR9 leads to NFkB activation and the induction of an immune response however, apical stimulation leads to the ubiquitination of IkBa which causes it to accumulate therefore blocking apical signaling (Lee et al., 2006). Together with the current work on TLR3, the asymmetric distribution of TLRs in IECs appears to represent a mechanism to avoid/control response against lumenal stimuli. Interestingly, a new study suggests that to achieve homeostasis, which we define here as co-habitation of IECs with the commensal flora, TLRs are differentially localized to specific sections in the GI tract (Price et al., 2018). Compatible to the present study, TLR2, 4, and 5 were found at both the apical and basolateral membranes. Interestingly, this study also showed that TLR7 and 9 were absent from the epithelium and their expression was limited to underlying immune cells. Our findings are consistent with this work as we also measured very low expression levels of TLR7/9 and observed that IECs are not responsive to the corresponding agonists. TLR3 was found to be expressed in all intestinal sections (Price et al., 2018, Pott et al., 2012, Cario and Podolsky, 2000). We show that TLR3 is specifically localized at the basolateral side of IECs providing these cells the possibility of mounting a polarized immune response. We could demonstrate that TLR3 displays a polarized functionality independently of its location within the GI tract. We found that basolateral infection leads to a stronger immune response in organoids derived from duodenum, ileum and colon (data not shown). Together these studies suggest that to achieve homeostasis, TLRs expression has to be spatially controlled both at the cellular level (apical/basolateral) and at the longitudinal level along the GI tract.

We report that AP-1B is responsible for targeting TLR3 to the basolateral side of IECs. The function of clathrin adapter proteins in participating in the regulation of TLR-mediated signaling was previously suggested. Adapter protein AP-3, has been shown to be critical for TLR4 and TLR7/9 mediated signaling (Mantegazza et al., 2012, Blasius et al., 2010). In the absence of AP-3, TLR7/9 are unable to produce type I IFN and TLR4’s recruitment and signaling from phagosomes is impaired. Additionally, AP-1 sigma1c mutations have been found in patients that exhibit a severe auto inflammatory skin disorder and it is suggested that this is due to misregulation of the TLR3 receptor further supporting our model of a key role for AP-1 in TLR3 signaling (Setta-Kaffetzi et al., 2014).

Due to the challenge associated with the commensal microbiota/epithelium interface, IECs finely tune their intrinsic immune system. Regulation can be found at the quantitative level by selecting the expression of a subset of TLRs or can be achieved by creating a uniquely tailored response that allows for directional partial tolerance of PAMPs. We propose that in addition to plasma membrane-associated TLRs, IECs also functionally polarize the intracellular endosomal TLR3. This polarization of function is intrinsically linked to the polarized nature of the cell and allows for TLR3 to strongly respond to basolateral challenges while limiting response to apical challenges (Sup. Figure 12). A recent study has shown that production of a type III interferon mediated antiviral immune response in mice was key to protect them from subsequent enteric virus infections (Ingle et al., 2019). This suggests that production of type III IFN against enteric viruses is key to protecting the gut against further challenges. In the context of our polarized immune response to viral infection, we propose that apical infection/challenges create a sufficient immune response allowing the non-infected bystander cells to be primed for further challenges. If this primed antiviral state is overruled by viruses, upon spreading and infection of IECs from their basolateral side, a stronger immune response will be generated acting as a second line of defense against the pathogen. Importantly, AP-1B knock-out mice show an increase in inflammation and infiltration of CD4+ T cells due to an overreaction of IECs against the commensal flora (Takahashi et al., 2011). In the line with our results that AP-1B functionally polarizes PRR sensing and signaling, we propose that within the physiological organization of the gut, this side specific polarized immune response of IECs participates in the tolerance of the commensal flora located in the lumen while remaining responsive to invading pathogens that will gain access to the sterile basolateral side.

## Acknowledgments

This work was supported by a research grant from Chica and Heinz Schaller Foundation and Deutsche Forschungsgemeinschaft (DFG) (Project number 240245660 (Project 14 of SFB 1129) and 278001972 (Project A09 of TRR186) to SB). This project has received funding from the European Union’s Seventh Framework Programme under grant agreement no 334336 (FP7-PEOPLE-2012-CIG). MS was supported by the Olympia Morata Fellowship from Heidelberg University Hospital, the Brigitte-Schlieben Lange Program from the state of Baden Württemberg, Germany and the Dual Career Support from CellNetworks, Heidelberg, Germany. MM is supported by the TRR186. KP and SM were supported by the SFB1129.

## Author Contributions

SB^a^=Steeve Boulant, SB^b^=Sina Bartfeld

MLS and SB^a^ designed experiments, MLS, SM, KP, MM and DA performed experiments, MLS analyzed data, TH analyzed microarray data, TK and HO generated the KO mice organoids, CO and JK assisted in establishing polarized infection. SB^b^ designed and performed microinjection, VH assisted for the polarized sorting, MLS and SB^a^ wrote the manuscript. The final version of the manuscript was approved by all authors.

## Materials and Methods

### Cell and Viruses

T84 human colon carcinoma cells (ATCC CCL-248) were maintained in a 50:50 mixture of Dulbecco’s modified Eagle’s medium (DMEM) and F12 (GibCo) supplemented with 10% fetal bovine serum and 1% penicillin/streptomycin (Gibco). Mammalian reovirus (MRV) strains Type 3 clone 9 (T3C9) derived from stocks originally obtained from Bernard N. Fields were grown and purified by standard protocols (Sturzenbecker et al., 1987).

### Antibodies and Inhibitors

Rabbit polyclonal antibody against MRV μNS was used at 1/1000 for immunostaining and western blots (Broering et al., 2000, Shah et al., 2017); ZO-1 (mouse from Santa Cruz Biotechnology and rabbit from Thermo Scientific) were both used at 1/100 for immunostaining; actin (Sigma-Aldrich) was used at 1/2000 for western blots; Phalloidin-AF647 (Molecular Probes) was used at 1/200 for immunostaining; Laminin (Abcam) was used at 1/100 for immunostaining, and TLR3 (Abcam) was used at 1/100 for immunostaining. Secondary antibodies were conjugated with AF488 (Molecular Probes), AF568 (Molecular Probes), AF647 (Molecular Probes), CW800 (Li-Cor) or HRP (Sigma-Aldrich) directed against the animal source. TRIF peptide inhibitor was used at a final concentration of 25 μM (Invivogen). Actinomycin D (Sigma) was used at a final concentration of 4 μg/mL. TLR stimuli: Flagellin-ST (InivoGen), ODN 2395, 2006, and 2243 (InvivoGen), LPS-EK (InvivoGen), poly I:C (Invivogen), poly I:C HMW Fluorescein (InvivoGen), Pam3CSK4 (InvivoGen) and R848 (InvivoGen) were used at the concentrations listed in figure legends.

### Viral infections

All MRV infections were performed at a MOI of 1. Media was removed from transwells and virus containing media was added to either the apical or basolateral side. Virus was maintained for the course of the experiment.

### Polarization of T84 cells on transwell inserts

1.2×10^5^ T84 cells were seeded on polycarbonate transwell inserts (Corning, polycarbonate, 3.0 μM) in DMEM/F12 medium. Medium was replaced 24h post-seeding and every two days subsequently. The trans-epithelial electrical resistance (TEER) was tested as indicated with EVOM^2^ apparatus (World Precision Instrument). When the TEER reached 1000 Ohm/cm^2^, the T84 cells were considered polarized (Madara et al., 1987). Polarization was controlled by immunostaining of the tight junction protein ZO-1 (see indirect immunofluorescence assay).

### Seeding of T84 cells into spheroids

T84 cells were dissociated using 0.25% trypsin, washed with PBS and resuspended in culture medium containing 4% Matrigel. Dissociated cells were seeded into Matrigel-coated eight-well chamber slides (Lab-Tek, eight-well chamber slides, Nunc, 155411) at a concentration of 12,000 single cells per well. Chamber slides were coated with 10 µl pure Matrigel and incubated at 37°C for 2 min before cell seeding. Cells were incubated for six days and fixed using 2% PFA for 20 min. Immunostaining of spheroids was carried out as described below.

### RNA isolation, cDNA, and qPCR

RNA was harvested from cells using NuceloSpin RNA extraction kit (Machery-Nagel) as per manufactures instructions. cDNA was made using iSCRIPT reverse transcriptase (BioRad) from 250 ng of total RNA as per manufactures instructions. q-RT-PCR was performed using SsoAdvanced SYBR green (BioRad) as per manufacturer’s instructions, TBP and HPRT1 were used as normalizing genes. For mouse interferon lambda and HPRT1 TaqMan gene expression assays were used.

### Indirect Immunofluorescence Assay

Polycarbonate transwell inserts were cut in half or cells were fixed in 2% paraformaldehyde (PFA) for 20 mins at room temperature (RT). Cells were washed and permeabilized in 0.5% Triton-X for 15 mins at RT. Primary antibodies were diluted in phosphate-buffered saline (PBS) and incubated for 1h at RT. Membranes were washed in 1X PBS three times and incubated with secondary antibodies for 45 mins at RT. Membranes were washed in 1X PBS three times and mounted on slides with ProLong Gold DAPI (Molecular Probes). Cells were imaged by epifluorescence on a Nikon Eclipse Ti-S (Nikon) or by confocal tile scans on a Zeiss LSM 780 (Zeiss). ZO-1 images were acquired on an ERS 6 spinning disc confocal microscope and deconvolution was performed using Huygens Remote Manager. Spheroids and 3D organoids were imaged on an inverted spinning disc confocal microscope (Nikon, PerkinElmer) with 60x (1.2 numerical aperture, PlanApo, Nikon) water immersion objective and an EMCCD camera (Hamamatsu C9100-23B).

### Western blot

At time of harvest, media was removed, cells were rinsed one time with 1X PBS and lysed with 1X RIPA (150 mM sodium chloride, 1.0% Triton X-100, 0.5% sodium deoxycholate, 0.1% sodium dodecyl sulphate (SDS), 50 mM Tris, pH 8.0 with phosphatase and protease inhibitors (Sigma-Aldrich)) for 5 mins at RT. Lysates were collected and equal protein amounts were separated by SDS-PAGE and blotted onto a nitrocellulose membrane by wet-blotting (Bio-Rad, Germany). Membranes were blocked with 5% milk or 5% BSA in TBS containing 0.1% Tween 20 (TBS-T) for one hour at room temperature. Primary antibodies were diluted in blocking buffer and incubated overnight at 4°C. Membranes were washed 3X in TBS-T for 5 mins at RT. Secondary antibodies were diluted in blocking buffer and incubated at RT for 1 h with rocking. For film exposure: membranes were washed 3X in TBS-T for 5 mins at RT. HRP detection reagent (GE Healthcare) was mixed 1:1 and incubated at RT for 5mins. Membranes were exposed to film and developed. For LiCOR assays, membranes were scanned on a LiCOR Odyssey imager.

### ELISA

Supernatants were collected at time points indicated in figure legends. Supernatants were kept undiluted. IFNλ2/3 was evaluated using the IFNλ2/3 DIY ELISA (PBL Interferon source) for the basolateral supernatants only as we have previously shown that IFNλ is preferentially secreted to the basolateral compartment (Stanifer et al., 2016). IL-6 was evaluated using the IL-6 DIY ELISA (Biolegend). ELISA was performed as per manufacturers instructions.

### Plaque Assay

BSC-1 were seeded into 24-well plates at a density of 200,000cells/well. 24-48h post-seeding when cells had reached 100% confluency, media was removed and cells were washed 1X with PBS+2mM MgCl_2_ (PBS-M). Virus samples were harvested from infected T84 cells and were subject to three rounds of freezing and thawing. Samples were spun to remove cellular debris and virus containing supernatants were diluted in 10-fold serial dilutions in PBS-M. Infection was allowed to proceed for 1h at RT with rocking every 15 mins. At the end of the incubation time, cells were overlaid with a 1:1 mixture of 2% agarose:2X 199 Media (Sigma-Aldrich) containing 10 μg/mL chymotrypsin. Cells were incubated at 37°C for 48h or until the appearance of plaques. Cells were fixed in 10% formaldehyde for 30 mins at RT. Plugs were removed and cells were stained with 0.5% crystal violet for 15 mins at RT. Crystal violet was removed and cells were washed with water. Plaques were counted and all samples were performed in triplicate.

### Fluorescent poly I:C uptake assay

1 x 10^5^ T84 cells were grown on collagen coated transwell filters until reaching full polarization. HMW poly I:C fluorescein was added to the apical or basolateral surface at a concentration of 10 μg/mL. After incubating at 37°C for the indicated time points, transwells were washed with PBS and then incubated with an acid wash (PBS, 0.2M acetic acid, 0.5M NaCl) for 10 sec to remove any non-internalized poly IC. Cells were then washed three times in 1X PBS followed by lysis in RIPA buffer. The amount of fluorescence of the internalized virus was measured on a Tecan infinite M200 Pro at 495 nm. Cells which were not fed poly IC or where poly IC was added and immediately removed were used as a background control.

### Dextran uptake assay

1 x 10^5^ T84 cells were grown on collagen coated transwell filters. At the indicated time, the diffusion of FITC-labelled dextran was measured. FITC-labeled dextran (Sigma-Aldrich, 4 kDa) was to the apical surface at a final concentration of 2 mg/mL. After incubating for 3 h 37°C, 100 µL aliquots of the basal media were collected and the fluorescence was measured with the FLUOstar Omega spectrofluorometer (BMG Labtech) at 495 nm. As positive control, fluorescence of 100 µL aliquot of a collagen coated but cell-free transwell filter was measured to assess maximum diffusion of FITC-labeled dextran.

### Fluorescent viral uptake assay

1 x 10^5^ T84 cells were grown on collagen coated transwell filters until reaching 1000Ω/cm^2^. AF488-labelled MRV (Boulant et al., 2013) was to the apical or basolateral surface at a MOI=1. After incubating at 37°C for the indicated time points, transwells were washed with PBS and then incubated with an acid wash (PBS, 0.2M acetic acid, 0.5M NaCl) for 10 sec to remove any non-internalized virus particle. Cells were then washed three times in 1X PBS followed by lysis in RIPA buffer. The amount of fluorescence of the internalized virus was then measured on a Tecan infinite M200 Pro at 495 nm. Cells which were not infected with virus or were virus was added and immediately removed were used as a background control.

### Human organoid cultures

Human tissue was received from colon and small intestine resection from the University Hospital Heidelberg. This study was carried out in accordance with the recommendations of the University hospital Heidelberg with written informed consent from all subjects in accordance with the Declaration of Helsinki. All samples were received and maintained in an anonymized manner. The protocol was approved by the “Ethics commission of the University Hospital Heidelberg” under the protocol S-443/2017. Stem cells containing crypts were isolated following 2 mM EDTA dissociation of tissue sample for 1 h at 4°C. Crypts were spun and washed in ice cold PBS. Fractions enriched in crypts were filtered with 70 μM filters and the fractions were observed under a light microscope. Fractions containing the highest number of crypts were pooled and spun again. The supernatant was removed and crypts were re-suspended in Matrigel. Crypts were passaged and maintained in basal and differentiation culture media (see table 1).

### Mouse organoids

Mouse intestinal tissue was received from control floxed mice or mice expressing a floxed Ap1m2 gene as previously described (Kanaya et al., 2018). Organoids were harvested, passaged and maintained in cultures as previously described for the human organoids, except using mouse specific media conditions (see table 1). To obtain organoids lacking the Ap1m2 gene, organoids were treated with 20 μM tamoxifen for 48h. Tamoxifen was removed and cells were allowed to recover and grow in normal mouse media.

### Microinjection

Human colon organoids were microinjected adapting a previously published protocol (Bartfeld and Clevers, 2015). Microinjection was carried out in a sterile laminar flow hood (KoJair, Finnland) using a micromanipulator (Narishige) and microinjector (Eppendorf FemtoJet). In each well, 30 organoids were injected. The injection needle was placed either inside the organoid (apical infection), or outside the organoid (basal infection), but both time in similar proximity to the epithelium. Pairs of inside and outside injections were on the same plate.

### Organoids on transwells

Tranwells were coated with 2.5% collagen in PBS for 1 h prior to organoids seeding. Organoids were collected at a ratio of 100 organoids/transwell. Collected organoids were spun at 450g for 5 mins and the supernatant was removed. Organoids were washed 1X with cold PBS and spun at 450g for 5 mins. PBS was removed and organoids were digested with 0.5% Trypsin-EDTA (Life technologies) for 5 mins at 37°C. Digestion was stopped by addition of serum containing medium. Organoids were spun at 450g for 5 mins and the supernatant was removed and organoids were re-suspended in normal growth media at a ratio of 100 μL media/well. The PBS/matrigel mixture was removed from the transwells and 100 μL of organoids were added to each well. 500 μL of normal organoid media was added to the bottom chambers. 24 hours post-seeding media on both sides of the transwells was changed for differentiation media and the TEER was monitored over five days.

### RNA decay assay

T84 cells were seeded onto a 24-well plate at a density of 100,000cells/well. 24h post-seeding T84 cells were infected with MRV T3C9. 16 hours post-infection, cells were treated with 4 μg/mL actinomycin D and RNA samples were collected at time points indicated in figure legend. RNA was harvested and qPCR as performed as described above.

### Production of Ap1 knock-down T84 cells

T84 cells were seeded onto 6-well plates in a density of 500,000cells/well. 24h post-seeding cells were transduced with lentiviruses expressing an shRNA targeted for the Ap1m2 gene (sequence available upon request). 72h post-transduction, lentiviruses were removed and T84 cells were put under antibiotic selection. When all control cells had died, antibiotic selection was removed and T84 cells were allowed to re-grow. Knock-down was confirmed through qPCR.

### Microarray

T84 cells were seeded on transwell and were allowed to polarize. Once a TEER value of 1000Ω/cm^2^ had been reached, cells were infected from the apical and basolateral membranes with MRV at a MOI=1. 16 hours post-infection RNA was collected and 1 μg of total RNA was used for an Illumina HumanHT-12 beadarray analysis. Expression values were quantile-normalized after background correction (Shi et al., 2010) and log2-transformed. Unspecific filtering was performed by discarding probes without significant detection p-value in any sample. Differentially expressed probes between A and Mock as well as between B and Mock were identified using the empirical Bayes approach (Smyth, 2004) based on moderated t-statistics as implemented in the Bioconductor package limma (Ritchie et al., 2015). P-values were adjusted for multiple testing using the Benjamini-Hochberg correction in order to control the false discovery rate. Adjusted p-values below 5% were considered statistically significant. For heatmap display, expression values of significantly up-regulated genes (in A or B) were scaled across samples and hierarchical clustering was performed using euclidean distance and complete linkage. Analyses were carried out using R 3.5 with add-on package pheatmap.

### Statistics

Statistical analysis was performed using the GraphPad Prism software package.

## Supplemental Information

**Supplementary Figure 1. Establishment of barrier function by T84 cells.** T84 cells were seeded onto transwell inserts. (A). The rate of establishment of barrier function was measured by the trans-epithelial electrical resistance (TEER) using a chop stick electrode. TEER greater than 1000 Ohm/cm^2^ indicates a tight barrier and is marked with a dashed line. (B). Barrier integrity was assessed by dextran diffusion assay. T84 cells were allowed to grow on transwells and at the indicated time points 4kDa fluorescent FITC-Dextran was added to the apical chamber. Three hours post-incubation, basolateral media was collected and analyzed by fluorometry to assess the amount of dextran which has diffused to the basolateral compartment of the transwells. Experiment were performed in triplicate. Error bars indicate the standard deviation. n.d.= not determined.

**Supplementary Figure 2. Activation of pBMC by TLR ligands and uptake of TLR3 agonist by T84 hIECs.** (A) pBMCs were isolated and treated with 10-fold serial dilutions of the indicated TLR agonists (starting concentrations: TLR1/2 (PAM3CSK4, 1 μg/mL); TLR3 (1:1 HMW and LMW poly I:C, 10 μg/mL); TLR4 (LPS-EK, 1 μg/mL); TLR5 (Flagellin-ST, 1 μg/mL); TLR7/8 (R848, 1 μg/mL), TLR9 (ODN 2395 5 μM)). Supernatants were harvested 24 hours post-stimulation and were tested for IL-6 by ELISA. (B) Polarized T84 cells were fed FITC-labeled poly I:C in a side specific manner. Cells were harvested at indicated time points, were washed with a quick acid wash to remove any non-internalized poly I:C bound to the cell surface. Following acid wash, cells were lysed and the relative amount of internalized FITC-labeled poly I:C was measured by spectrofluorometry. Experiments were performed in triplicate. Error bars indicate the standard deviation. ns=not significant (unpaired t-test).

**Supplementary Figure 3. Apical infection of T84 cells by MRV leads to more *de novo* virus production compared to basolateral infection.** Polarized T84 cells were infected apically or basolaterally with MRV. (A) Fluorescently labeled MRV was added to polarized T84 cells in a side specific manner. Cells were collected at indicated times points and were washed with a quick acid wash to remove any non-internalized virus particles bound to the cell surface. Following acid wash, cells were lysed and the relative amount of internalized virus particles was measured by spectrofluorometry. (B) Infected cells were collected at the indicated time points and viral replication was assayed by q-RT-PCR against the MRV μ2 genome segment. (C) Same as B except virus replication was addressed by Western blot against the MRV non-structural protein μNS. Actin was used as a loading control (left panel). Representative figure is shown. The relative expression of μNS is normalized to actin is shown (right panel). (D) Infected cells were fixed at indicated time points and immunofluorescence was performed against the non-structural protein μNS. The number of infected cells per field of view was counted. 10 fields of view were counted for each time point. (E) Polarized T84 cells were infected apically or basolaterally with MRV. Infected cells were collected at the indicated time points and viral replication was assayed by q-RT-PCR against the MRV μ2 genome segment. Experiments were performed in triplicate, error bars indicate the standard deviation. ns=not significant, *<P.05, **P < 0.01 (unpaired t-test)

**Supplementary Figure 4. Side specific induction of IFN production upon viral infection in the human intestinal epithelial line SKCO15 cells.** SKCO15 cells were seeded on transwells. Following establishment of barrier function, cells were infected apically or basolaterally with MRV. (A) 16 hpi, the production of type I (IFNβ) and type III (IFNλ2/3) IFNs was assessed by q-RT-PCR. (B) Efficiency of the primary MRV infection in SKCO15 cells was controlled by immunofluorescence against the non-structural protein μNS (red). Cell nuclei were stained using DAPI (blue). Experiments were performed in triplicate. Representative images for IF are shown. Scale bar =100 μm. Error bars indicate standard deviation. ** P < 0.01 (unpaired t-test).

**Supplementary Figure 5. Basolateral infection of T84 cells leads to a stronger production of interferon compared to apical infection.** (A) T84 cells were infected apically or basolaterally with MRV. RNA samples were collected at the indicated times post-infection and production of type I (IFNβ) and type III (IFNλ2/3) interferons was measured by q-RT-PCR. (B) T84 cells were infected apically or basolaterally with MRV. Supernatants were collected at the indicated times post-infection and analyzed by ELISA for the secretion of type III (IFNλ2/3) interferon. A=apical infection. B =basolateral infection. Black line indicates the limit of detection. All samples were performed in triplicate. Error bars indicate the standard deviation. ns=not significant, *<P.05, **P < 0.01, ***P < 0.001 (unpaired t-test).

**Supplementary Figure 6. Stronger immune response observed during basolateral infection of hIECs is not due to differences in transcript stability.** (A) Polarized T84 cells were infected apically or basolaterally with MRV. 16 hpi cells were treated with actinomycin D to block transcription. RNA samples were collected at the indicated times post-actinomycin treatment. (B) Relative amounts of type I (upper panel) and type III IFN (lower panel) transcripts (normalized to the time of actinomycin treatment) were evaluated by q-RT-PCR (left panel). The half-life of each transcripts was determined (right panel). Experiments were performed in triplicate. Error bars indicate the standard deviation. ns=not significant. *<P.05 (unpaired t-test).

**Supplementary Figure 7. Basolateral infection of T84 cells by MRV leads to the upregulation of many immune related genes.** Polarized T84 cells were infected apically or basolaterally with MRV. 16 hpi, RNA was collected and subjected to microarray analysis. Expression values were quantile-normalized after background correction and log2-transformed. Unspecific filtering was performed by discarding genes with non-significant p-values in any sample. (A) Differentially expressed genes between apical infection and mock infected cells as well as between basolateral infection and mock infected cells were identified using the empirical Bayes approach and are shown in a Venn diagram. (B) Heatmap of expression values of significantly up-regulated genes in apically and basolaterally infected cells were scaled across samples and hierarchical clustering was performed using euclidean distance and complete linkage. (C) KEGG analysis of significantly upregulated genes upon basolateral infection of T84 cells.

**Supplementary Figure 8. Basolateral infection of human organoids by MRV generates a stronger intrinsic immune response compared to apical infection.** (A) Schematic of the microinjection approach used to infect organoids specifically from either their apical or basolateral sides. (B-C) Human intestinal organoids were infected by MRV apically or basolaterally through microinjection. Virus infection was assessed 16 hpi for (B) the production of the non-structural viral protein μNS using Western blot analysis and (C) the production of the MRV μ2 genome segment by q-RT-PCR. (D) Generation of intrinsic immune response was assessed by quantifying the production of type I (IFNβ) and type III (IFNλ2/3) IFNs using q-RT-PCR. (E) The relative expression of TLR2-9 was evaluated in human intestinal organoids. Results are expressed as a ratio to the house keeping gene HPRT1. All experiments were performed in triplicate. Representative western blot is shown. Error bars indicate the standard deviation. ns=not significant, *<P.05, **P < 0.01 (unpaired t-test).

**Supplementary Figure 9. Knock-down of the clathrin adaptor AP-1B impacts the polarized immune response.** (A) Knock-down efficiency AP-1B was evaluated by q-RT-PCR. (B-D) The integrity of the barrier function of the AP-1B knock down T84 cells was evaluated by (B) indirect immunofluorescence of the tight junction protein Zo-1 (red), (C) dextran diffusion assay and (D) TEER measurement.(E) Polarized WT and AP-1B knock-down T84 cells were infected with MRV in a side specific manner. Supernatants were collected at indicated time points and subjected to ELISA to monitor the amount of secreted type III IFNs. All experiments were performed in triplicate. Error bars indicate the standard deviation. Representative image for immunofluorescence is shown. Scale bar =100 μm. ns=not significant, ** P < 0.01, **** P < 0.0001 (unpaired t-test).

**Supplementary Figure 10. Stimulation of AP-1B knock-down T84 cells by TLR agonists.** (A) AP-1B knock-down T84 cells were seeded on transwells and treated with indicated TLR agonists: TLR1/2 (PAM3CSK4, 1 μg/mL); TLR3 (1:1 HMW and LMW poly I:C, 10 μg/mL); TLR4 (LPS-EK, 0.01 μg/mL); TLR5 (Flagellin-ST, 1 μg/mL); TLR7/8 (R848, 1 μg/mL), TLR9 (ODN 2395 5 μM). 6 hours post-treatment RNA was harvested and evaluated for the upregulation of IL-6 by q-RT-PCR. Results are normalized to mock-treated samples. (B) Same as A except type III IFN (IFNλ2/3) was evaluated. Experiments were performed in triplicate; error bars indicate the standard deviation. ns=not significant, *<P.05 (unpaired t-test)

**Supplementary Figure 11. Inhibition of TRIF interferes with the polarized immune response generated by T84 cells.** WT and AP-1B knock-down T84 cells were seeded on transwell inserts. (A) WT cells were treated with a TRIF inhibitor or a control peptide. T84 cells were infected with MRV in a side specific manner. RNA samples were harvested at indicated time points post-MRV infection and the intrinsic innate immune induction was analyzed by q-RT-PCR for type III IFN (IFNλ2/3). (B) Same as A, except AP-1B knock-down cells were evaluated. All experiments were performed in triplicate. Error bars indicate the standard deviation. ns=not significant, * P < 0.05. ** P < 0.01. Statistics in panel B are between AP-1B and AP-1B+TRIFi.

**Supplementary Figure 12. The clathrin sorting adaptor protein AP-1 localizes TLR3 to the basolateral side of hIECs leading to an asymmetric IFN-mediated immune response.** Schematic showing TLR3 localization in polarized intestinal epithelial cells and its implication for side specific interferon induction upon viral infection.

